# Microbial composition of the human nasopharynx varies according to influenza virus type and vaccination status

**DOI:** 10.1101/647693

**Authors:** Tao Ding, Timothy Song, Bin Zhou, Adam Geber, Yixuan Ma, Lingdi Zhang, Michelle Volk, Shashi N. Kapadia, Stephen G. Jenkins, Mirella Salvatore, Elodie Ghedin

## Abstract

Factors that contribute to enhanced susceptibility to severe bacterial disease after influenza infection are not well defined, but likely include the microbiome of the respiratory tract. Vaccination against influenza, while having variable effectiveness, could also play a role in microbial community stability. We collected nasopharyngeal samples from 215 individuals infected with influenza A/H3N2 or influenza B and profiled the microbiota by target sequencing of the 16S rRNA gene. We identified signature taxonomic groups by performing linear discriminant analysis and effective size comparisons (LEfSe) and defined bacterial community types using Dirichlet Multinomial Mixture (DMM) models. Influenza infection was shown to be significantly associated with microbial composition of the nasopharynx according to the virus type and the vaccination status of the patient. We identified 4 microbial community types across the combined cohort of influenza patients and healthy individuals with one community type most representative of the influenza-infected group. We also identified microbial taxa for which relative abundance was significantly higher in the unvaccinated elderly group; these taxa include species known to be associated with pneumonia.

**Importance:** Our results suggest that there is a significant association between the composition of the microbiota in the nasopharynx and the influenza virus type causing the infection. We observe that vaccination status, especially in more senior individuals, also has an association with the microbial community profile. This indicates that vaccination against influenza, even when ineffective to prevent disease, could play a role in controlling secondary bacterial complications.

## Introduction

Influenza virus is the major cause of severe viral respiratory infection in adults, resulting in more than 200,000 hospitalizations and 30,000 to 50,000 deaths each year in the US (1). A critical factor in influenza virus-associated morbidity and mortality is the increased susceptibility of infected individuals to bacterial pneumonia, a common complication of influenza pandemics (2, 3) and epidemics (4). Epidemiological studies have shown that despite circulating in humans since 1968, seasonal H3N2 influenza outbreaks are associated with increased clinical severity, including excess respiratory mortality and excess pneumonia and influenza hospitalizations (5). During the 2014-2015 season, the antigenically drifted H3N2 influenza virus caused major outbreaks globally, resulting in increased pneumonia- and influenza-associated mortality (http://www.cdc.gov/flu/weekly/weeklyarchives2014-2015/week2.htm#S2) (6).

Although annual trivalent influenza vaccines are available and widely received, vaccine effectiveness can be limited. Vaccines are less effective in the elderly, a population that is particularly vulnerable to influenza infections and that tends to develop more severe influenza complications (7). In general, H3N2 influenza strains have been associated with lower antibody responses and decreased vaccine effectiveness even in well-matched years (8, 9). During the 2014-2015 influenza season, vaccine effectiveness in the Northern Hemisphere against H3-specific influenza was estimated to be at 22% (95% CI, 5–35%) (10). This low effectiveness was attributed both to the low immunogenicity of the H3 vaccine components (11), and to a mismatch of the H3 component with the circulating H3 viruses (6). Despite the limitations of the 2014-2015 influenza vaccine to prevent disease caused by these antigenically drifted strains, a study carried out in the hospital setting suggested that vaccination could have prevented a more severe disease requiring hospitalization (12).

The reasons for enhanced susceptibility to severe bacterial disease after influenza infection remain poorly defined. Bacteria like *Staphylococcus aureus* and *Streptococcus pneumoniae*, which are the most prominent pathogens involved in bacterial super-infection, are common colonizers of the upper respiratory tract (URT) and make up the URT microbiome with other resident microbes. However, viral infection can disrupt this equilibrium and cause loss of some microbial populations and/or overgrowth of other pathogens, resulting in disease. Disruption of the URT microbiota has been found to be associated with community acquired pneumonia (13), although it is still unclear whether the microbial changes observed are the cause or the consequence of the viral infection. The protective role of resident microbes has also been studied. For example, *S. aureus* priming mediates recruitment of M2 alveolar macrophages, which reduce influenza pathogenesis by limiting inflammation in the lungs (14).

One aspect not fully explored is how influenza types and strains impact the microbiota in the respiratory tract and whether vaccination could be protective by either reshaping the microbiota or preventing the virus from disrupting its equilibrium. In this regard, even an unmatched influenza vaccine could prevent severe disease by modulating the respiratory bacterial communities. Recent studies have shown an effect of the live attenuated influenza vaccine (LAIV) on the microbiota of the nasopharynx (15, 16). Although the microbiota of the respiratory tract has been described as the gatekeeper of respiratory health (17), information on how it changes in influenza infection, and how it could be impacted by vaccination is still sparse. To address these questions, we characterized the microbiota of the nasopharynx by analyzing samples from individuals diagnosed with Influenza A virus (IAV) H3N2 or Influenza B virus (IBV) (Yamagata or Victoria) infections during the 2014-2015 influenza season in New York. Investigating in more detail the relationship between host factors such as age or vaccination status, and the respiratory microbiome in a specific influenza season could help us better identify factors that contribute to influenza disease severity.

## Results

We analyzed a total of 226 NP swabs from 215 patients diagnosed with IAV (n=157) or IBV (n=58) collected at New York Presbyterian hospital/Weill Cornell Medicine in New York City during the 2014-2015 influenza season. One patient (patient ID 213 in **Table S1**) was diagnosed with IAV first, and then with IBV a month later. Clinical characteristics of individual subjects and influenza vaccination history, including whether the patient had been vaccinated in previous seasons, in the current season (2014-2015), or both, is summarized in **Table S1**. Of individuals infected with IAV, 43% (67/157) had been vaccinated in the current season while 62% (97/157) had received the influenza vaccine in one or more of the 5 previous seasons (**Table 1**). Of individuals infected with IBV, 41% (24/59) were vaccinated in the current season and 53% (31/59) at some point in the last 5 seasons (**Table 2**).

**Table 1.** Characteristics of subjects with influenza A included in the study.

| **Patient characteristics** | **Overall**<br>**n (%)** | **Young**<br>**(<18 yo)**<br>**n (%)** | **Adult**<br>**(18-64 yo)**<br>**n (%)** | **Elderly**<br>**≥65 n (%)** |
| --- | --- | --- | --- | --- |
| Total number | 157 | 31 | 60 | 66 |
| Age, median (IQR) | 60 (24-77) | 3 (0-7) | 46 (33-55) | 79 (72-86) |
| Male gender | 60 (40%) | 15 (48%) | 20 (33%) | 25 (38%) |
| **Care Setting** |  |  |  |  |
| ER | 32 (20%) | 14 (45%) | 12(60%) | 6 (9%) |
| Inpatient | 71 (45%) | 8 (26%) | 17 (28%) | 46 (70%) |
| Outpatient/Clinic | 54 (34%) | 9 (29%) | 31 (52%) | 14 (21%) |
| **Immunocompromise<sup>1</sup>** |  |  |  |  |
| Yes | 40 (25%) | 4 (13%) | 19 (32%) | 17 (26%) |
| No | 104 (66%) | 24 (77%) | 34 (57%) | 46 (70%) |
| Unknown | 13 (8%) | 3 (10%) | 7 (12%) | 3 (5%) |
| **Documented LRTI<sup>2</sup>** |  |  |  |  |
| Yes | 15 (10%) | 1 (3%) | 4 (7%) | 10 (15%) |
| No | 126 (80%) | 26 (84%) | 47 (78%) | 53 (80%) |
| Unknown | 16 (10%) | 4 (13%) | 9 (14%) | 3 (5%) |
| **Vaccination current season**<br>**(2014-2015)** |  |  |  |  |
| Yes | 67 (43%) | 11 (35%) | 23 (38%) | 33 (50%) |
| No | 64 (41%) | 18 (58%) | 30 (50%) | 16 (24%) |
| Unknown | 26 (16%) | 2 (6%) | 7 (12%) | 17 (26%) |
| **Vaccination past 5 years** |  |  |  |  |
| Yes | 97 (62%) | 14 (45%) | 38 (63%) | 45 (68%) |
| No | 45 (29%) | 16 (52%) | 15 (25%) | 14 (21%) |
| Unknown | 15 (9%) | 1 (3%) | 7 (12%) | 7 (11%) |
| **Disposition / Outcome** |  |  |  |  |
| Home | 135 (86%) | 28 (90%) | 49 (82%) | 58 (88%) |
| Hospitalized | 5 (3%) | 0 | 2 (3%) | 3 (5%) |
| Death | 3 (2%) | 0 | 0 | 3 (5%) |
| Unknown | 14 (9%) | 3 (10%) | 9 (15%) | 2 (3%) |

Abbreviations: IQR = interquartile range, ED = emergency department, LRTI = lower respiratory tract infection
<sup>1</sup> Includes neutropenia, leukemia, lymphoma, stem cell transplant, solid organ transplant, pregnancy, HIV, use of high dose steroids, monoclonal antibodies, or systemic chemotherapy.
<sup>2</sup>Diagnosed by chest x-ray

**Table 2.** Characteristics of subjects with influenza B included in the study

| **Patient characteristics** | **Overall**<br>**n (%)** | **Young**<br>**(<18 yo)**<br>**n (%)** | **Adult**<br>**(18-64 yo)**<br>**n (%)** | **Elderly**<br>**(≥65 yo)**<br>**n (%)** |
| --- | --- | --- | --- | --- |
| Total number | 58 | 8 | 28 | 22 |
| Age, median (IQR) | 59 (42-71) | 2.5 (1-8) | 51 (43-61) | 75(69-88) |
| Male gender | 30 (51%) | 4 (57%) | 15 (50%) | 10 (50%) |
| **Care Setting** |  |  |  |  |
| ED | 8 (14%) | 1 (12%) | 4 (14%) | 3 (14%) |
| Inpatient | 20 (34%) | 3 (38%) | 3 (11%) | 14 (64%) |
| Outpatient | 30 (52%) | 4 (50%) | 21 (75%) | 5 (23%) |
| **Immunocompromise<sup>1</sup>** |  |  |  |  |
| Yes | 10 (17%) | 1 (12%) | 6 (21%) | 3 (14%) |
| No | 42 (72%) | 7 (87%) | 18 (64%) | 17 (77%) |
| Unknown | 6 (10%) | 0 | 4 (14%) | 2 (9%) |
| **Documented LRTI<sup>2</sup>** |  |  |  |  |
| Yes | 6 (10%) | 0 | 1 (4%) | 5 (21%) |
| No | 51 (88%) | 8 (100%) | 24 (92%) | 19 (79%) |
| Unknown | 1 (2%) | 0 | 1(4%) | 0 |
| **Vaccination current season**<br>**(2014-2015)** |  |  |  |  |
| Yes | 24 (41%) | 4 (50%) | 7 (25%) | 13 (59%) |
| No | 13 (22%) | 4 (50%) | 7 (25%) | 2 (9%) |
| Unknown | 21 (36%) | 0 | 14 (50%) | 7 (32%) |
| **Vaccination past 5 years<sup>3</sup>** |  |  |  |  |
| Yes | 31 (53%) | 5 (63%) | 7 (25%) | 19 (86%) |
| No | 7 (12%) | 2 (25%) | 5 (18%) | 0 |
| Unknown | 20 (34%) | 1 (12%) | 16 (57%) | 3 (14%) |
| **Disposition / Outcome** |  |  |  |  |
| Home | 57 (98%) | 8 (100%) | 28 (100%) | 21 (95%) |
| Death | 1 (2%) | 0 | 0 | 1 (5%) |

Abbreviations: IQR = interquartile range, ED = emergency department, LRTI = lower respiratory tract infection
<sup>1</sup>Includes neutropenia, leukemia, lymphoma, stem cell transplant, solid organ transplant, pregnancy, HIV, use of high dose steroids, monoclonal antibodies, or systemic chemotherapy.
<sup>2</sup>Diagnosed by chest x ray
<sup>3</sup>Children too young to receive flu vaccine (< 1y) are not counted

### The microbiota of the nasopharynx is significantly different in influenza-infected subjects compared to uninfected individuals

To determine whether the microbial community of the nasopharynx was different in the context of influenza infection, we compared NP swabs collected from influenza-infected patients to those from 40 healthy individuals (controls). Although no significant difference in microbial alpha-diversity was detected between the influenza and control groups (Shannon index and inverse Simpson index were tested, p < 0.05), the microbial compositions were significantly different (AMOVA test: p-value < 0.001) leading to samples clustering within each group, as visualized by multidimensional scaling (**Fig. 1**). We compared the relative abundance and prevalence of taxa that were represented in both groups, i.e. the core microbiota, (**Fig. 2a**), and identified signature taxa for each group using linear discriminant analysis and effective size comparisons (LEfSe) analysis **(Fig. 2b)**. The control samples appeared to be dominated with *Corynebacterium* and *Streptococcus*, while the influenza-infected individuals had a slightly lower prevalence and abundance of *Streptococcus*, an enrichment of *Dolosigranulum*, and very low prevalence of *Corynebacterium* (**Fig. 2a**).

**Figure 1.**
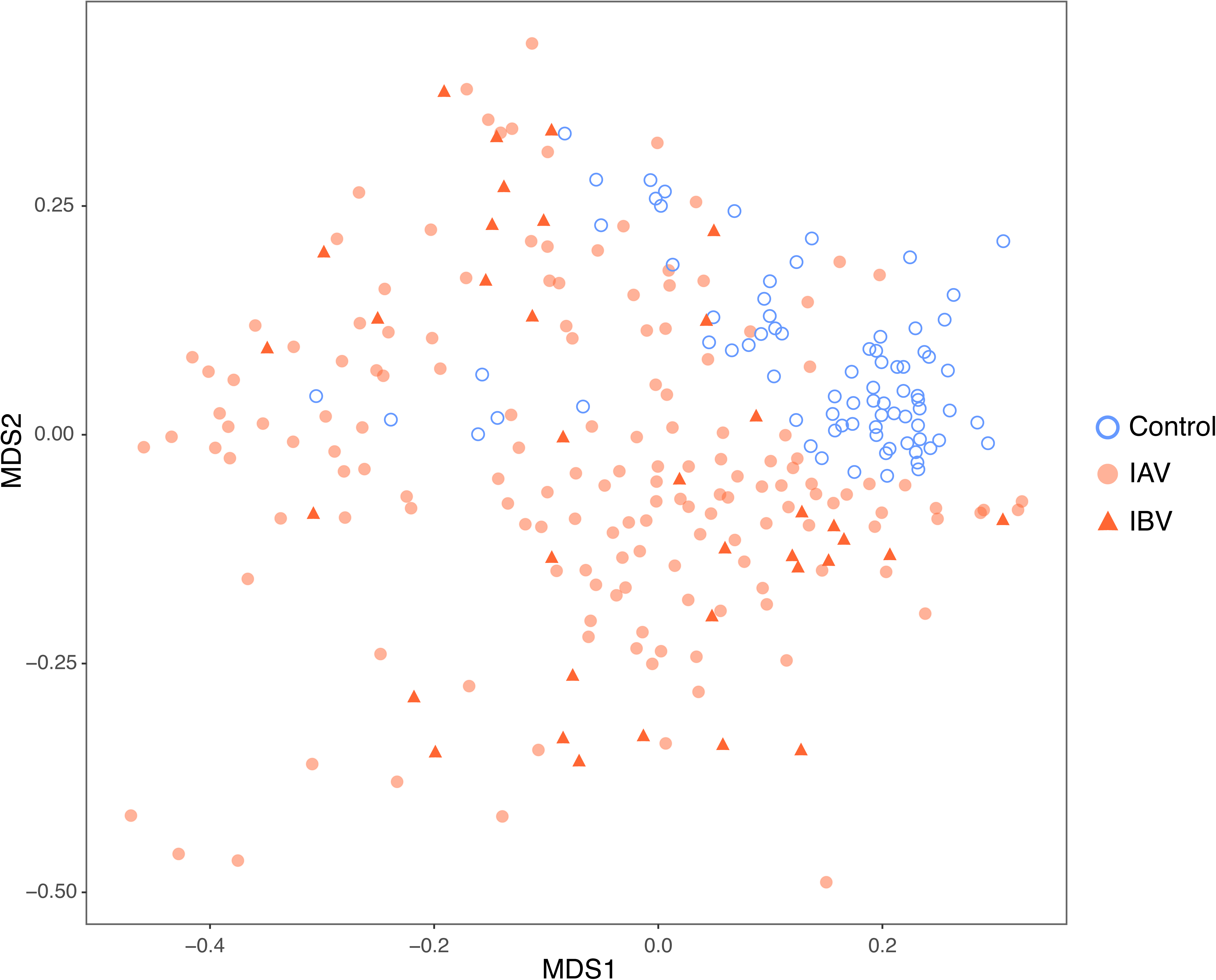
Clustering of influenza-infected samples and healthy control samples based on genus-level taxonomic assignments. Clustering is displayed as a NMDS plot of all the samples, in which the dissimilarity between samples is calculated as the Bray-Curtis distance.

**Figure 2.**
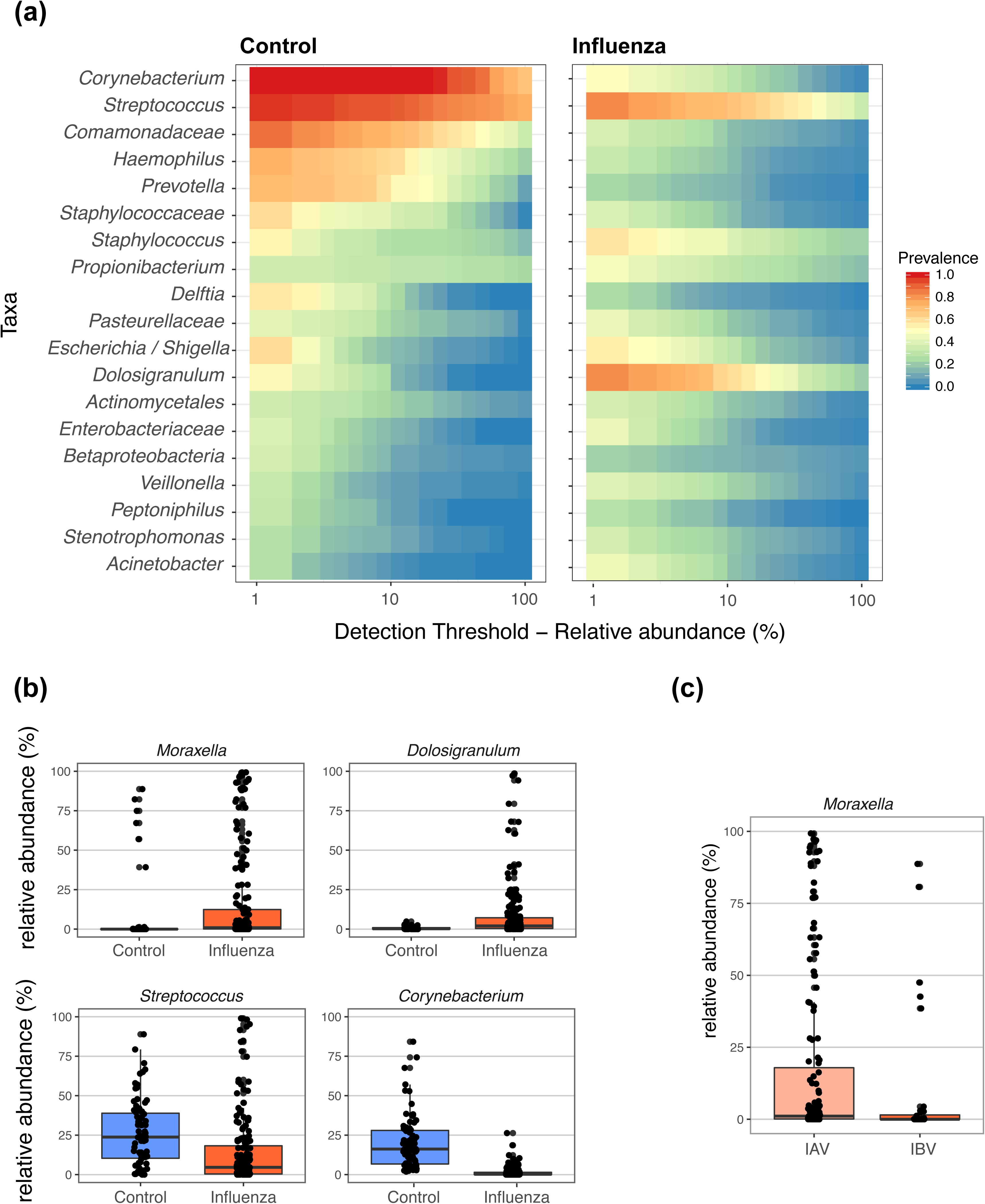
**(a)** Core microbiota heatmaps showing abundance of taxa and prevalence across samples in healthy controls and influenza-infected samples. Taxa listed were selected based on their prevalence in the two groups of samples. **(b)** Relative abundance of significant taxa enriched in influenza infection (top graphs) or in healthy controls (bottom graphs). Significance was determined by LEfSe. Whiskers represent values outside the upper and lower quartiles**. (c)** Relative abundance of significant taxa enriched in influenza A virus infection as compared to influenza B virus infection. Significance was determined by LEfSe. Whiskers represent values outside the upper and lower quartiles.

We also used LEfSe to identify taxonomic features that were most likely to significantly characterize the influenza-related compositional differences of the microbiota. We observed 4 taxa (**Table S2**), including *Moraxella* and *Dolosigranulum*, that were significantly enriched in the influenza group (**Fig. 2b**, *upper panel*). We identified 7 key taxa (**Table S2**), with the top 2 being *Streptococcus* and *Corynebacterium*, which had a relative abundance that was significantly higher in the control group (**Fig. 2b**, *lower panel*).

Considering that IBV is often associated with milder disease and to determine if IBV infection was associated with a different microbial profile than IAV, we compared the NP microbiota of IAV- and IBV-infected subjects. Moderate differences in their compositions were detected (**Fig 1**; p.value < 0.01 by AMOVA). Using LEfSe we observed that the relative abundance of 4 taxa, including *Moraxella* (**Fig. 2c**) were significantly higher in IAV infections (**Table S3**) while 4 taxa were significantly higher in IBV infections (**Table S3**).

We further explored the association between influenza infection and microbial community structure. To do so we partitioned the data into community types using Dirichlet multinomial mixture models. We identified 4 microbial community types (NP-types) in the subjects tested. NP-type A was significantly enriched in influenza infection while NP-type B dominated in the control group (**Fig. 3a; Table 3**). While both community types C and D were slightly enriched in influenza patients, it was not at a significant level. To study the effect of age on the core microbiome of influenza-infected subjects, we divided them in 3 groups: young (<18 years), adult (18-64 years) and elderly (65 year or older). In the influenza-infected subjects only, the less common influenza-associated NP-type D was found predominantly in the young (<18 yo), (**Fig. 3b**), while NP-type A was under-represented in this group. We analyzed the abundance of the dominant taxa in each NP-type (**Fig. 3c**). NP-type A was comprised primarily of *Streptococcus* and *Dolosigranulum*; NP-type B, enriched in the control cohort, was dominated by a combination of *Streptococcus, Corynebacterium*, and *Comamonadaceae*. NP-type C was dominated by *Moraxella* while NP-type D had an overrepresentation of *Staphylococcus*. The different communities did not associate differently with either of the flu types (IAV vs IBV).

**Table 3.** Number of samples associated with each community type.

|  | NP-type A | NP-type B | NP-type C | NP-type D |
| --- | --- | --- | --- | --- |
| Healthy | 2 | 59 | 4 | 14 |
| Infected | 105 | 3 | 20 | 63 |

**Figure 3.**
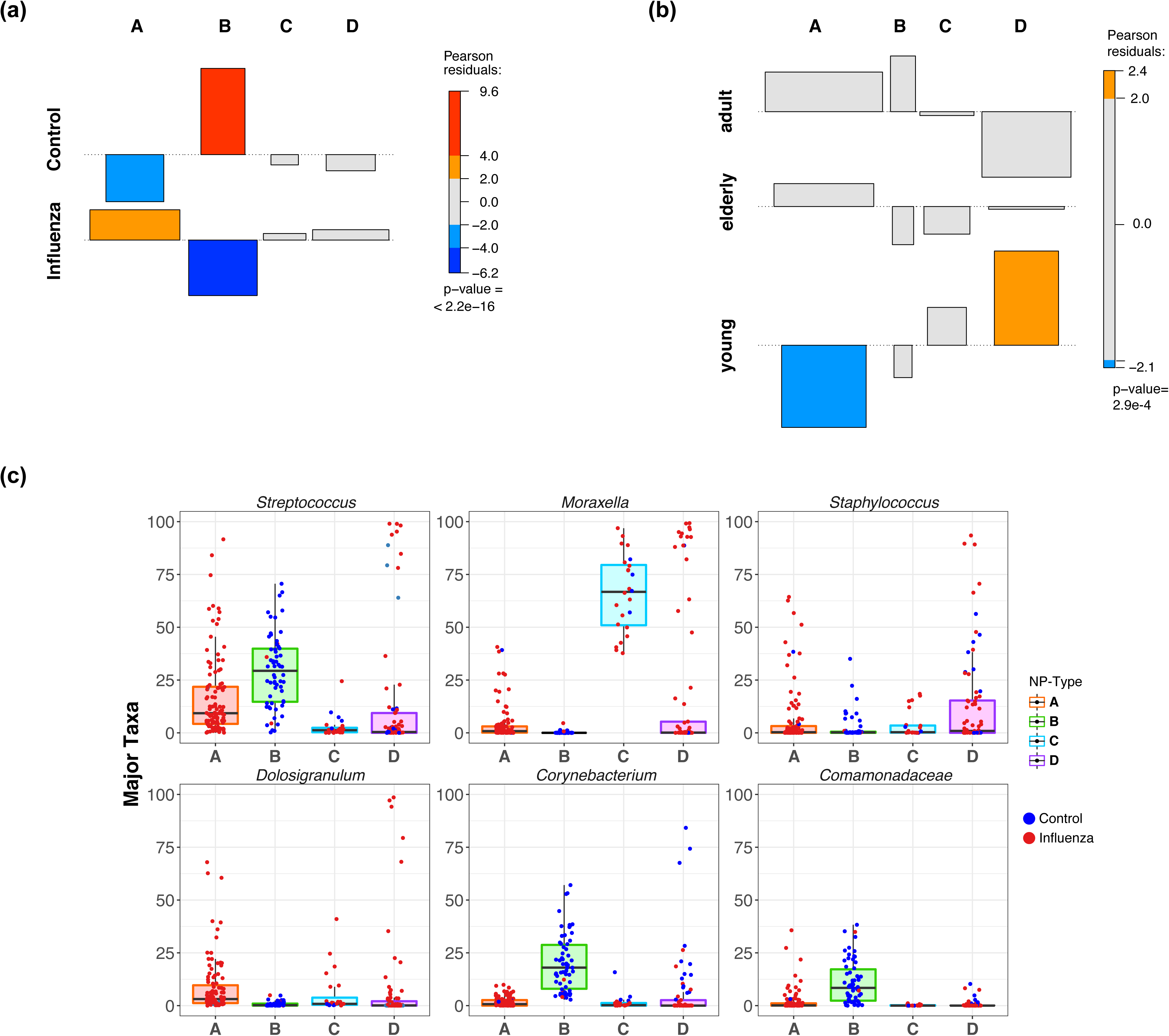
**(a)** Association between the four NP-types and influenza infection status, determined by Chi-square test. **(b)** Association between the four NP-types and the three age groups in influenza-infected subjects. **(c)** Relative abundance of the dominant microbial taxa in each NP-type.

### Association of vaccination with the nasopharynx microbiota is different in IAV and IBV

We determined whether vaccination in the current season had an association with the microbial composition of the NP in the influenza-infected individuals. We did not observe differences in sample clustering between individuals who were vaccinated or not, indicating that microbial composition was similar in both groups (**Fig 4a**; p.value >0.01 in AMOVA). However, when looking for taxonomic features that significantly characterized each group, we observe by LEfSe analysis an enrichment of specific taxa in the unvaccinated subjects—*Moraxella* in IAV (p=0.002), and *Streptococcus* in IBV (p=0.010) (**Fig. 4b)**.

**Figure 4.**
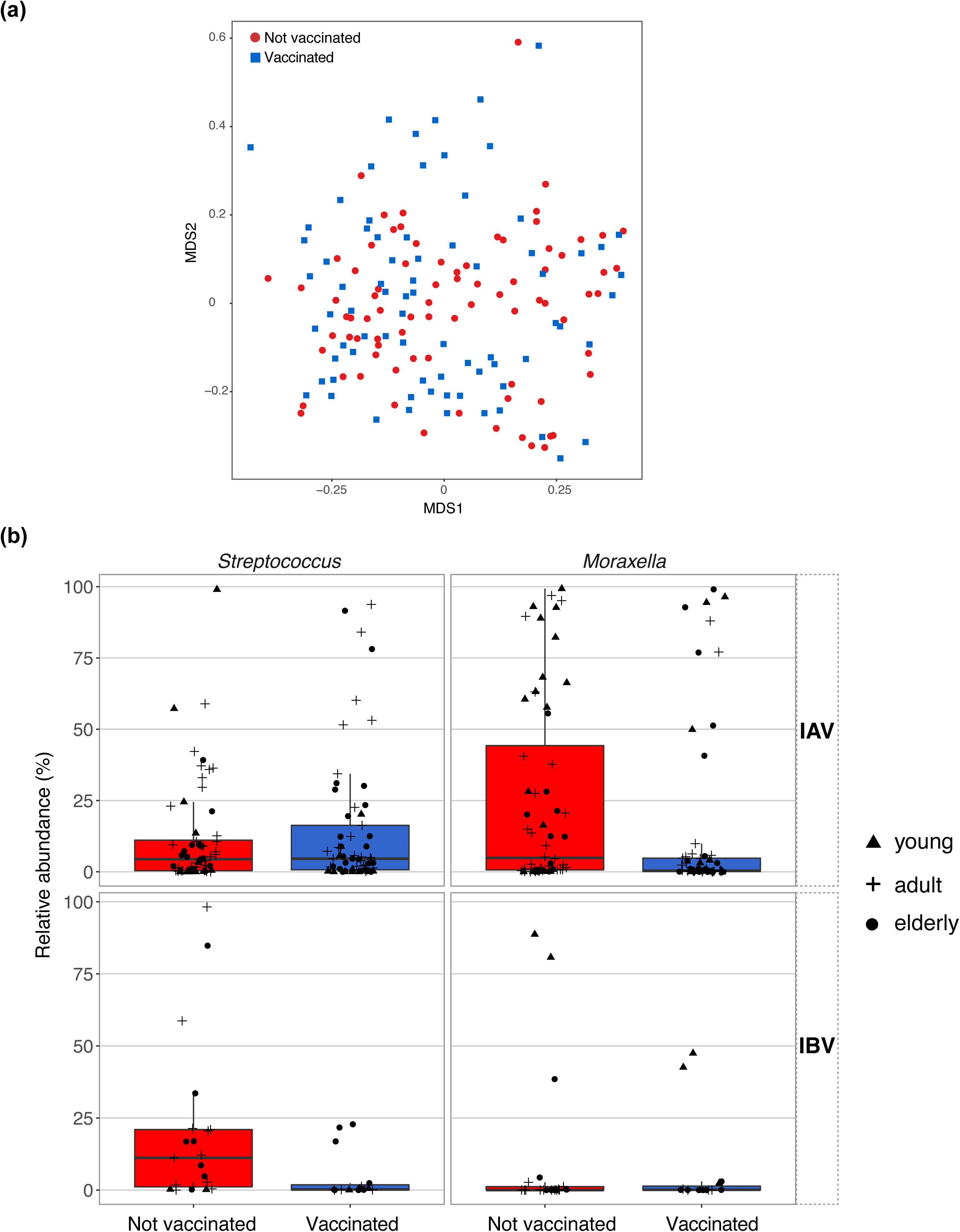
**(a)** Beta diversity ordination calculated by NMDS using the Bray-Curtis dissimilarity of samples from vaccinated and unvaccinated individuals. **(b)** Relative abundance of *Streptococcus* and *Moraxella* in unvaccinated patients (left) and vaccinated patients (right), separated by IAV (top) and IBV infections (bottom). Age group of the individual from which sample was collected is indicated by different symbols.

We also tested for age-dependent differences in the microbiota of the nasopharynx in vaccinated and unvaccinated individuals infected with either IAV or IBV, but we did not observe any significant differences in enriched taxa between age groups. However, when testing overall microbial diversity (as measured by Shannon entropy), we observe higher microbial diversity in the unvaccinated elderly than in the vaccinated elderly (Wilcox test p.value = 0.005) **(Fig. 5a**); we did not observe a similar effect in the two other age groups. To further study what compositional differences contributed to this age-specific difference in microbial diversity, we identified with LEfSe 7 microbial taxa for which relative abundance was significantly higher in the unvaccinated elderly group (**Fig. 5b)**. Some members of these taxa such as Staphylococcaceae, Gram-negative bacteria (Pasteurellaceae and Escherichia/Shigella), and Sphingomonas also include species known to be associated with post-influenza (including pneumonia) and nosocomial infections (18). Finally, we tested if any of the clinical variables (listed in **Table 1** and **Table S1**), including pneumonia, antibiotic usage, Tamiflu usage, and immunocompromise, was associated with specific features of the nasopharyngeal microbiota, but we did not find any significant association.

**Figure 5.**
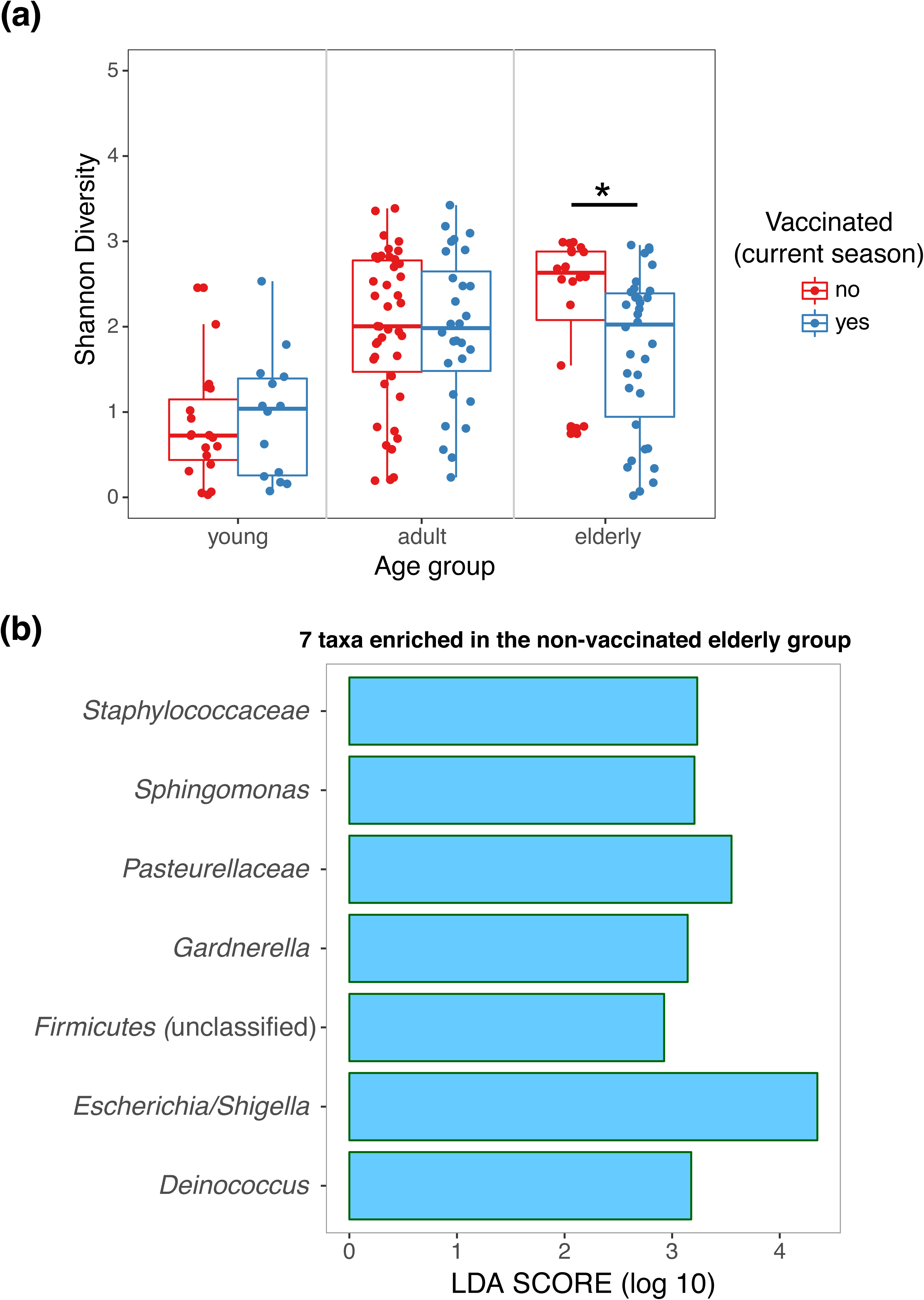
**(a)** Comparison of alpha diversities (calculated as Shannon Index) of unvaccinated patients with those of vaccinated patients, separated by age groups. **(b)** Linear discriminant analysis (LDA) score of the seven microbes found to be significantly enriched in unvaccinated elderly patients against vaccinated elderly, determined by LEfSe.

### Influenza genetic diversity affects the microbiota of the nasopharynx

While we observed differences in microbial enrichment between individuals infected with IAV and IBV, and in community types for infected versus controls, we explored whether there was also an association between IAV genetic diversity and the microbial community within the nasopharynx. We first performed a K-mer analysis to identify underlying influenza sequence signatures for each sample and compared them to each other, visualizing this measure of genetic distance by multidimensional scaling (**Fig. 6a)**. Three clusters were identified for influenza A/H3N2, with 2 clusters corresponding to the 3C.2a genetic clade. While we did not see a correspondence between the sample clustering profile and the NP-type microbial profiles (data not shown), we did observe by performing LEfSe analysis that one of the two 3C.2a clusters, HA-2, had a significantly higher relative abundance of *Escherichia (Shigella)* (**Fig. 6b**) as compared to the other 3C.2a cluster (HA-3) and the 3C.3 cluster (HA-1). While all 3 clusters had samples with high relative abundance of *Staphylococcus*, it was significantly different across the 3 HA groups, with potentially higher relative abundance in the group corresponding to clade 3C.3 (HA-1).

**Figure 6.**
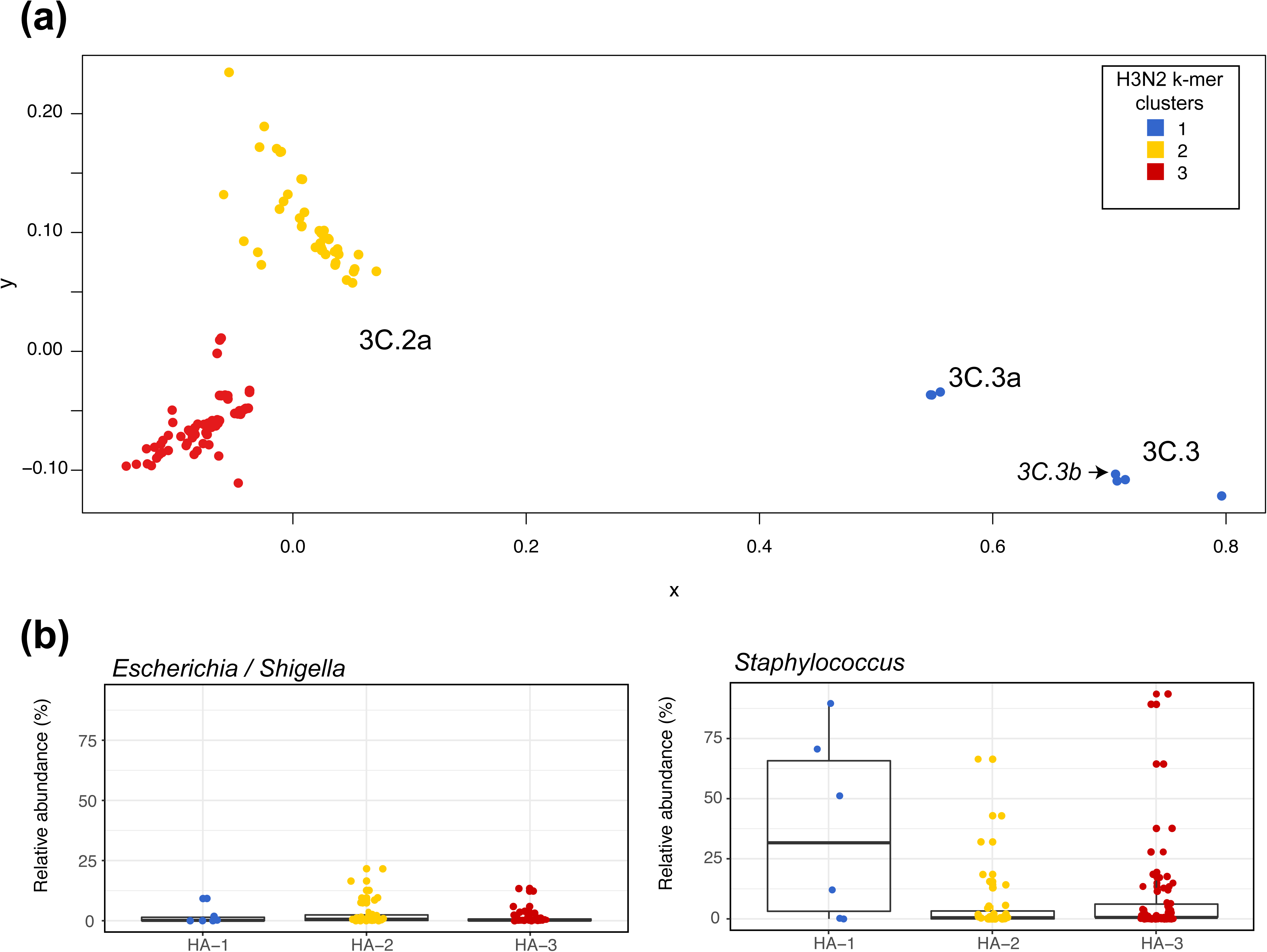
**(a)** Clustering of IAV patients based on the genetic diversity of the HA segment. **(b)** Relative abundance of the microbes that vary across the HA genetic clusters.

## Discussion

A number of studies have looked at the respiratory tract microbiota and influenza infections (15, 19-21), but this is the first to explore the respiratory microbiota in both IAV H3N2 and IBV infection in the context of vaccination in a year with low vaccine effectiveness. One study characterized the microbiota in patients infected with the 2009 pandemic H1N1 influenza. They used a *cpn60* amplicon sequencing method and no healthy control was involved, so no inference was made regarding the alteration of the microbiota caused by influenza infection (19). We have shown that IAV and IBV virus infections were associated with a significantly different microbial community profile than uninfected individuals. The microbiota of the nasopharynx in infected individuals was enriched with taxa such as *Dolosigranulum* and *Staphylococcus* when compared to the microbiota of the control group. These two genera have previously been shown to be associated with pneumonia (22, 23). *Dolosigranulum* are Gram-positive bacteria for which currently only one species has been identified (*D. pigrum)*. Although a rare opportunistic pathogen, *D. pigrum* has been confirmed as a causative agent in different types of pneumonia and septicemia (22, 23). The significant enrichment of these types of Gram-positive bacteria as compared to healthy controls indicates that the predisposition to super-infection could be initiated early on in the infection, likely due to in part to the dysregulation of the innate and immune response by the virus (reviewed in (23)). We also see a significantly higher relative abundance of *Moraxella spp*—Gram-negative bacteria—in influenza-infected subjects as compared to the healthy control group. A number of species in this genus are resident microbes of mucosal surfaces, occasionally leading to opportunistic infections. *Moraxella spp*. have already been recognized as human respiratory tract pathogens (24) and seen in some cases to lead to influenza infection complications (25). Interestingly an increased prevalence of *Moraxella spp.* has been reported in individuals with acute viral upper respiratory infections caused by viruses (26).

In the analysis of community types, we saw that one type in particular (NP-type A) was enriched in influenza-infected subjects and was comprised primarily of *Dolosigranulum* and *Streptococcus*. We also identified a negative association between influenza infection and *Corynebacterium*. *Corynebacterium* has been found to commonly colonize the human nose and skin and was shown to be overrepresented in children free of *Streptococcus pneumoniae* (27). *Corynebacterium accolens* was shown to inhibit the growth of *S. pneumoniae* by releasing antibacterial free fatty acids (27). A negative correlation between *S. aureus* and *Corynebacterium* abundance was also previously observed (28) with a recent study showing that *Corynebacterium* inhibits the virulence of *S. aureus* (29). In our own data, we also observed that *Staphylococcus* was present at very low relative abundance in NP-type B that was enriched in the control group and dominated by a combination of *Streptococcus, Corynebacterium*, and *Comamonadaceae*. Because species-level resolution for taxonomic assignment is difficult with 16S rRNA gene sequence analysis, we do not know whether *S. pneumoniae* was the dominant species for the *Streptococcus* identified in both NP-type A and NP-Type B. Overall, these observations suggest that *Corynebacterium* could potentially protect the respiratory tract from pathogenic bacteria such as *S. aureus* and *S. pneumoniae* that are the most common cause of post-influenza pneumonia. More work is needed to confirm that the observed lack of *Corynebacterium* in the nasopharynx of influenza-infected individuals contributes to the increased likelihood of influenza-induced pneumonia.

Since our study was cross-sectional and we studied the composition of the nasopharynx microbiota at the moment of diagnosis, we cannot determine whether differences between healthy controls and influenza-infected individuals are due to the infection, due to the presence of a microbial community that predisposes to infection, or a combination of both. A longitudinal study of the sputum from rhinovirus-infected individuals showed a rise in bacterial burden with a higher prevalence of *Haemophilus influenzae* associated with infection (30). Longitudinal influenza infection studies have reported conflicting results with one showing that the administration of live-attenuated influenza virus (LAIV) can modify the microbiota of the nasal cavity (16), while a study where volunteers challenged with an H3N2 strain were sampled over a 30-day period did not show any changes in the oropharyngeal microbiota (21). This lack of an effect may be due to the fact that the cohort was comprised of young and healthy volunteers with many who developed very mild disease (19 out the 52 challenged individuals). In contrast, our cohort includes patients from all age groups, including young and elderly, with a range of disease severity. A recent household transmission study shows that influenza susceptibility is associated with differences in the overall bacterial community structure, with a particularly increased influenza risk in young children (20). These differences between studies suggest that patient characteristics such as age, comorbidities, vaccination status and treatments and viral characteristics need to be considered when studying the effect of influenza infection on the respiratory microbiota.

Our study also addresses the association of vaccination with differences in the microbiota of the nasopharynx during influenza infection. The effectiveness of the influenza vaccine varies in different seasons (31, 32) due to a number of factors, such as vaccine strain mismatch and host immune status, including history of previous influenza vaccination. We suggest that another potential factor is the host microbiome. Recent studies have shown that the human microbiota, by impacting immune cell development and differentiation, could influence adjuvant and vaccine efficacy (33). LAIV was shown to affect the microbiota of the nasopharynx (15, 16), and lead to an increased abundance of specific microbes associated with IgA responses (15). A study on the effects of trivalent LAIV on bacterial carriage in the nasopharynx of toddlers showed that there was an increase in *S. pneumonia* and *M. catarrhalis* density 28 days after vaccination (34), indicating that the influenza virus, even when attenuated, could impact carriage density.

We show that in influenza-infected individuals the lack of vaccination in the current season is associated with the enrichment of different microbial taxa, such as *Moraxella* and *Streptococcus*, depending on the type of influenza virus (IAV vs IBV). Although we cannot exclude or confirm that other confounding factors may also play a role in shaping the nasopharyngeal microbiota of influenza-infected patients, we tested for factors that were included in our demographic data, such as age, sex, and antibiotic usage, and did not observe any other significant association. However, an aspect missing is how vaccination can specifically reduce the risk for respiratory comorbidities, which can be largely attributed to the disruption of the microbial community within the respiratory tract (17). Because of an increased risk of infection, young and elderly populations get the most benefit from influenza vaccination (35). We found that in the elderly group (65+), the microbial diversity in the nasopharynx of unvaccinated patients was significantly higher than in the vaccinated, with an overrepresentation of taxa that include pathogenic species associated with nosocomial infections. A recent study has linked increased nasopharynx microbial diversity with pneumonia infections in the elderly population (13). While even an unmatched influenza vaccine can provide some level of cross-protection, our findings suggest that a protective effect could also be mediated by modifications in the microbiota that can help limit the growth of opportunistic pathogens.

## Conclusion

Our aim in this study was to determine whether the microbiota of the nasopharynx was different in individuals with influenza infection, and identify factors associated with the variations observed between infected subjects in different age groups. We found that during influenza infection, the nasopharyngeal microbiota of vaccinated individuals was strongly associated with higher levels of specific microbial taxa, with different microbial profiles relative to virus types and clade. These observations provide new insight into influenza infection and highlight a need for more studies to explore the mechanism of how influenza vaccines—live-attenuated or killed—interact with the respiratory microbiota.

## Materials and Methods

### Subjects and sample acquisition

Nasopharyngeal (NP) swabs collected from subjects of any age and sex that were sent to the New York Presbyterian Hospital microbiology laboratory for influenza testing in the 2014-2015 season were used for this study. All samples were confirmed by Film Array (Biofire) to be either IAV H3N2 or IBV positive. Clinical data were abstracted from the electronic medical record. For every subject we collected data on patient demographics, comorbidities and related treatments, influenza vaccination history, underlying malignancy status and treatments, antibiotics and antiviral treatments, clinical course including infections and therapies, and microbiology data. We also collected 80 NP swabs from 40 healthy patients living in New York City as controls. These were enrolled as part of an IRB-approved study aiming to characterize the respiratory microbiome in immunocompromised patients and healthy controls (*manuscript in preparation*) (**Table S4**). These volunteers represented a mix of hospital clinic workers and community members as we wanted to establish whether the hospital environment contributed to the microbiota observed since a number of our patients were hospitalized. We did not observe any difference between hospital workers and community members. Total DNA and RNA were extracted from each sample and subjected to 16S rRNA gene sequencing for microbiota profiling and influenza virus gene segment sequencing, respectively.

### DNA extraction and 16S rRNA gene sequencing

DNA extractions from the NP swab specimens were performed using the PowerSoil DNA Isolation Kit (MO BIO Laboratories Inc) in a sterilized Class II Type A2 Biological Safety Level 2 Cabinet (Labgard ES Air, NuAire). The swabs were processed in batches, and the cotton tip of each swab was cut off and transferred into the PowerBead tubes as the starting material. Nuclease Free Water (Ambion, ThermoFisher Scientific Inc) was also processed through the same DNA extraction procedure as the specimens and the healthy individual specimen control. Extracted DNA was eluted in 50 µl nuclease-free water and stored at –20 °C until processing. Extracted DNA was then used in a PCR reaction to amplify the V4 hypervariable region of the 16S rRNA gene using primer pair 515F/806R to prepare the sequencing library (36). Six µl of extracted DNA from swab samples were used as template in a final volume of 25 µl, with 0.35 µl Q5 Hot Start High-Fidelity DNA Polymerase (New England BioLabs Inc), 5 µl 5X Q5 Buffer, 0.5 µl dNTP mix, and 0.5 µM forward and reverse primers. Thermal cycling conditions were 94 °C for 2 min, then 33 cycles of 94 °C for 30 s, 55 °C for 30 s, and 72 °C for 90 s, followed by 72 °C for 10 min. PCR products were purified using 0.65× volumes of AMPure XP Beads (Beckman Coulter) and eluted into 20 µl low TE (10mM Tris, 0.1mM EDTA), pH 8.0 on the Bravo Automated Liquid Handling Platform (Agilent Technologies). Eluted PCR products were quantified with a Quant-iT double-stranded DNA (ds-DNA) High-Sensitivity Assay Kit (Invitrogen) on an Infinite M200 Plate Reader (Tecan) according to the manufacturer’s instructions and were combined with equal input mass into a sequencing pool. The pool was then purified again with 0.65X volumes of AMPure XP Beads and analyzed on a 2200 TapeStation (Agilent Technologies) using a High Sensitivity D1000 ScreenTape (Agilent Technologies) to confirm the integrity of the sequencing library. Finally, the sequencing pool was quantified by qPCR using the KAPA Library Quantification Kit (KAPA Biosystems) on a Roche 480 LightCycler. The library was sequenced at the Genomics Core Facility of the Center for Genomics and Systems Biology, New York University using an Illumina PE 2×250 V2 kit on an Illumina MiSeq Sequencer.

### 16S rRNA gene sequence analysis

The sequencing data was processed using the 16S rRNA gene sequence curation pipeline that was implemented in the mothur software package (37) following a previously described procedure (38). Briefly, the raw sequences as fastq files were extracted from sff files, and any sequence that had mismatches to the barcode, more than one mismatch to the primers, more than 8 nucleotide homopolymers, or ambiguous base calls was removed. Trimmed sequences were de-noised using PyroNoise (39) and then aligned against a customized SILVA database (40). Chimaeric sequences were detected and removed using a de novo Uchime algorithm that was implemented in mothur (41). The De-chimaeric sequences were classified using the naïve Bayesian Classifier trained against a customized version of the RDP training set (v9). A minimum classification score of 80% was required and 1,000 pseudo-bootstrap iterations were used. The taxonomy of the remaining sequences was used to assign the sequences to genus-level phylotypes, also known as operational taxonomic units (OTU), and this allowed us to make a table of counts for the number of times each phylotype was observed in each sample. Phylotypes that were identified in less than 20% of the total samples were removed from subsequent analysis. Samples with fewer than 1,000 reads were removed from downstream analysis and all samples were sub-sampled or rarified to 1,000 reads to perform subsequent analyses. Signature microbial groups were identified by performing LEfSe (Linear discriminant analysis effective size) analysis (41) implemented in mothur. Bacterial community types were defined using a Dirichlet Multinomial Mixture (DMM) algorithm based method that was previously described and implemented in mothur (42). Statistical tests, including Wilcoxon signed-rank test, chi-square test, and student t test were performed in R.

### RNA extraction and viral segment sequencing

Total RNA was extracted from each sample according to the manufacturer’s recommendations using 100 µL of Viral Transport Media as input for the RNeasy Micro Kit (Qiagen). Influenza genomic RNA was subsequently converted into cDNA and amplified (40 cycles of PCR) via the SuperScript III One-Step RT-PCR System with Platinum Taq High-Fidelity DNA Polymerase (Invitrogen) according to previously published methods (43, 44). Each successfully amplified influenza RNA sample was prepared for sequencing by one of two methods. Concurrent experimental work confirmed that samples prepared with both methods yielded identical minor variant profiles. Sixty-five of the samples used for subsequent analyses were sonicated in a microTUBE (Covaris) using the S220 Focused-ultrasonicator (Covaris). The fragmented cDNA was purified by 0.8× volumes AMPure XP beads on the Agilent Bravo and quantified via the Quant-iT High-Sensitivity dsDNA Assay Kit. Fifty ng of cDNA from each sample were used an input for the NEBNext Ultra DNA Library Prep Kit for Illumina (New England Biolabs) according to the manufacturer’s recommendations. The remaining cDNA samples were prepared for sequencing using a modified version of the Nextera DNA Library Preparation Kit protocol (Illumina). Amplicons were purified by 0.8× volumes AMPure XP beads on the Agilent Bravo and quantified via the Quant-iT High-Sensitivity dsDNA Assay Kit before normalization to constant concentration (2.5 ng/µl); 2.5ng of cDNA from each sample were used as input for Nextera library preparation. Individual libraries prepared by either method were quantified via the Quant-iT High-Sensitivity dsDNA Assay Kit and pooled with equal input mass before re-purification and size-adjustment with 0.6x volumes AMPure XP beads. Each of the three resultant pools (one prepared with NEBNext, two prepared with Nextera) was quantified by qPCR using the KAPA Library Quantification Kit on a Roche 480 LightCycler and its size distribution was measured on a 2200 TapeStation using a D1000 ScreenTape (Agilent Technologies). Each pool was sequenced at the Genomics Core Facility at the Center for Genomics and Systems Biology, New York University using an Illumina PE 2×250 V2 kit on an Illumina MiSeq Sequencer. Each pool was seeded at 12pM and included a 10% PhiX spike-in to compensate for potential low base diversity.

### Viral sequencing data analysis

Samples were trimmed using trimmomatic and the trimmed reads of each sample were mapped using the Burrows-Wheeler Alignment Tool (bwa) (45) with default parameters against the A/New York/03/2015 H3N2 strain for IAV H3N2 infected samples and against both B/Kentucky/28/2015 (Victoria) and B/New York/WC-LVD-15-007/2015 (Yamagata) strains for IBV infected samples. These were then processed for quality filtering and analysis with samtools (46). The average quality of any given read had to pass a phred score of 25. For generation of consensus sequences, any given site had to be covered by at least 200 reads. Minority variants were discovered by using statistical tests to minimize false positive single nucleotide variant (SNV) calls that can be caused by sequence specific errors. This involves using a binomial test to ensure that reads come from both the forward and reverse orientation. Additional thresholds for minority variant detection include a frequency of 1%, which accounts for the sequencing noise as determined by a plasmid control sample, and a coverage of 500×.

## List of abbreviations

IAV: Influenza A Virus

IBV: Influenza B Virus

SNV: Single Nucleotide Variants

NP: Nasopharyngeal

URT: Upper Respiratory Tract

16S rRNA gene: 16S subunit of the ribosomal RNA gene

OTU: Operational Taxonomic Unit

QIIME: Quantitative Insights into Microbial Ecology

DMM: Dirichlet Multinomial Mixture

LEfSe: Linear Discriminant Analysis Effect Size

PCA: Principal component analysis

NMDS: Non-Metric Multidimensional Scaling

LDA: Linear Discriminant Analysis

## Declarations

### Ethics approval

The study was approved by the Weill Cornell Medical College Institutional Review Board and the New York University Institutional Review Board (Weill Cornell Medicine IRB protocol #1506016280).

### Consent for publication

Not applicable

### Availability of data and material

The sequencing data from this study are available in the Sequence Read Archive (SRA) under the following accession numbers: SRP132207 for 16S rRNA genes and PRJNA431639 for influenza virus data.

### Author Contributions

TD analyzed the sequence data, performed the statistical analyses, interpreted data, and wrote the manuscript; MS collected samples; AG, YM, LZ and MV did sample processing, sequence library preparation, and data analysis; TS performed influenza sequence data analysis; MS and EG developed the concept, supervised all analyses and interpretation; all authors approved the manuscript. BZ designed the influenza sequencing strategy; SNK extracted patient’s clinical and laboratory data; SJ performed sample collection and flu diagnosis.

## Acknowledgements

We thank the Genomics Core Facility of the Center for Genomics and Systems Biology at New York University, and the clinical staff at Weill Cornell School of Medicine.

## SUPPLEMENTAL MATERIAL

**Table S1.** Clinical and sequencing information of single samples included in the study.

**Table S2.** Taxa identified by LefSe as significantly enriched in influenza-infected or control group.

**Table S3.** Taxa identified by LefSe as significantly enriched in influenza A- or influenza B-infected subjects.

**Table S4.** Characteristics of healthy volunteers who provided samples.

